# Hidden impacts of environmental stressors on freshwater: Cytoscape® correlation networks for community ecotoxicology analysis

**DOI:** 10.1101/834622

**Authors:** V.L. Lozano

**Affiliations:** Universidad de Buenos Aires, Facultad de Ciencias Exactas y Naturales, Depto. Ecología, Genética y Evolución, Buenos Aires, Argentina; CONICET – Universidad de Buenos Aires. Instituto de Ecología, Genética y Evolución de Buenos Aires (IEGEBA), Universidad de Buenos Aires, Buenos Aires, Argentina

**Keywords:** Stressors, Freshwater, Community, Correlation networks

## Abstract

Freshwater systems are affected by multiple stressors. Ecotoxicological studies using communities or higher ecological levels often are focussed on abundance analysis. We suggest incorporating correlation networks to bring to light possible hidden impacts on community interactions.

## Introduction

Freshwater systems are exposed to multiple stressors such as pesticides, microplastics, heavy metals, wastewater, industrial residues, biological invasions, nutrification and physical factors such as a temperature increase due to global warming (Jackson et al. 2016).

Aquatic ecosystems are composed by several communities that could be direct or indirectly affected and interact through trophic relationships (Kattwinkel et al. 2010). The impact of stressors on freshwater communities can be detected by structural changes (specific abundances, diversity indexes, etc.) and/or by physiological effects on organisms, leading or not, to functional changes (productivity, gas emissions, etc.) (Havens, 1994). Running correlation networks using an interface as Citoscape® (Shannon et al. 2003) could help us understand the impact of a stressor on the relationships between the different components of a community (Lozano et al. 2019), and elucidate ecological mechanisms of mitigation or exacerbation. A new mode of response to a stressor at the community-level -or higher-could arise by studying the correlations of the changes in the abundances of the different components of a system. In this work, a community-based correlation analysis is proposed using specific abundances to assess possible hidden effects of environmental stressors with focus on freshwater systems.

## Methods

Simulated values of specific abundances in a community were used to assess possible hidden effects of one environment stressor. Spearman correlation coefficients were calculated with InfoStat v.2008 and correlation networks were built with Cytoscape® v.3.7.1. incorporating only significant correlation coefficients (p□0.001). Other simulated examples were built directly on Cytoscape® to illustrate proposals.

## Results and discussion

Simulated values of specific and total abundances of a community treated with a stressor are shown in Table I, with means and standard deviations of 4 replicates of control (C) and treatment (T). Non-statistical differences are seen under a classical abundance analysis (Fig. I).

**Table I:** Simulated values of specific abundances of control (C) and treated systems (T).

|  | Sp.1 | Sp.2 | Sp.3 | Sp.4 | Total |
| --- | --- | --- | --- | --- | --- |
| **C1** | 4 | 4 | 8 | 3 | 19 |
| **C2** | 2 | 2 | 4 | 1 | 9 |
| **C3** | 2 | 2 | 4 | 1 | 9 |
| **C4** | 2 | 2 | 4 | 1 | 9 |
| **Mean** | 2.5 | 2.5 | 5.0 | 1.5 | 11.5 |
| **SD** | 1.0 | 1.0 | 2.0 | 1.0 | 5.0 |
| **T1** | 2 | 4 | 4 | 1 | 11 |
| **T2** | 2 | 2 | 8 | 1 | 13 |
| **T3** | 4 | 2 | 4 | 3 | 13 |
| **T4** | 2 | 2 | 4 | 1 | 9 |
| **Mean** | 2.5 | 2.5 | 5.0 | 1.5 | 11.5 |
| **SD** | 1.0 | 1.0 | 2.0 | 1.0 | 1.9 |

**Fig. I:**
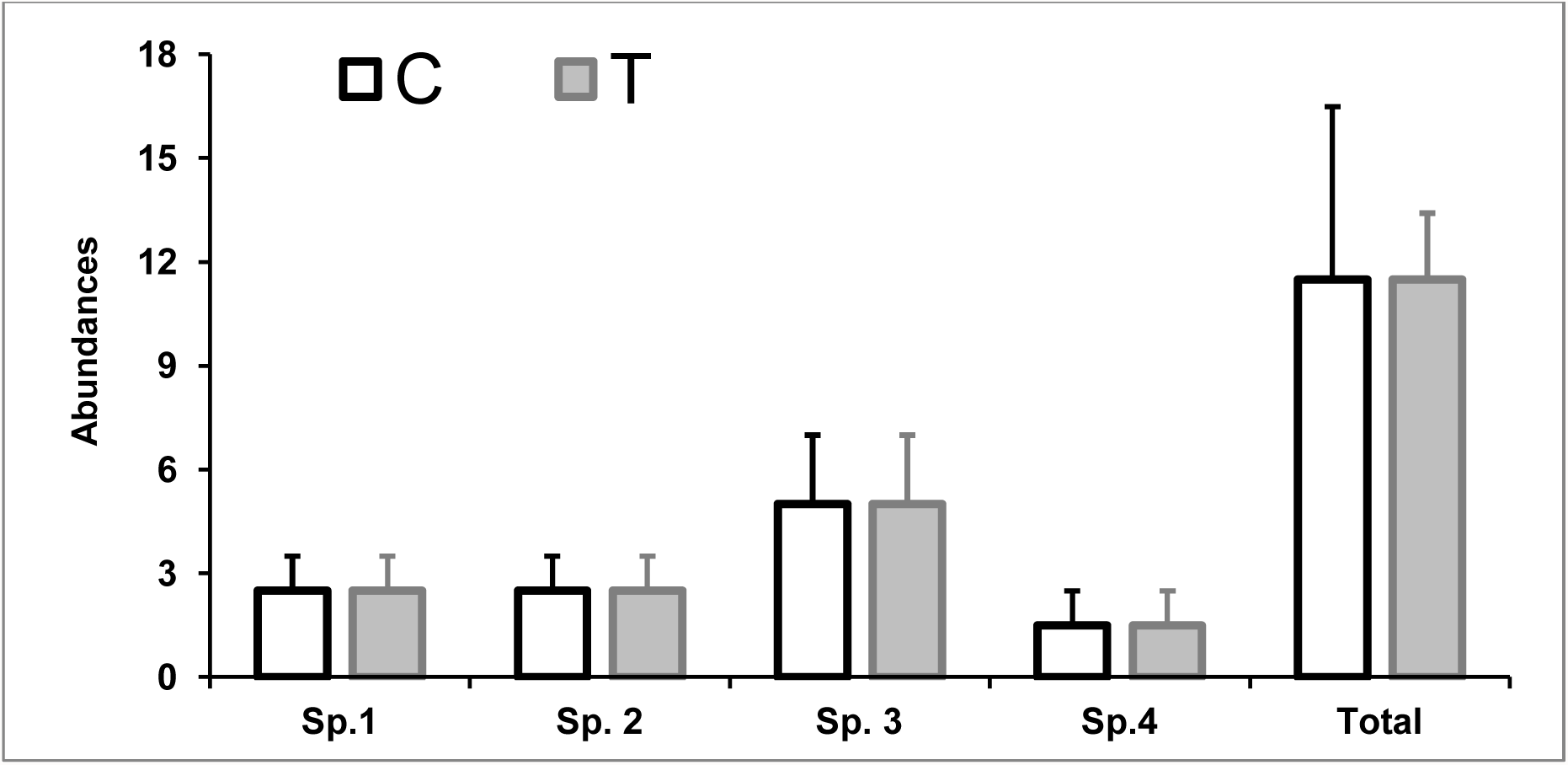
Abundances of simulated control (C) and treated (T) systems, non-significant variations are found.

Using the specific abundances of Table I, Spearman correlations were calculated and used to build networks with Cytoscape®. The hidden effects of the stressor changing the connectivity between the populations are shown (Fig. II). Considering the number of replicates (n=4) and the number of specific abundances (4), the maximum possible correlations were 6 (4*3:2) and the minimum was 0 if any specific abundance was correlated. Results show that a community with non-significant variations of specific abundances could be strongly affected by the modifications of its internal connectivity between species. In this example, while in both situations (A and B) specific abundances of each species show no effect, correlations between species shift from a situation where all populations from a community are positively correlated (A) to a situation where each one is totally disconnected from any other (B).

**Fig. II:**
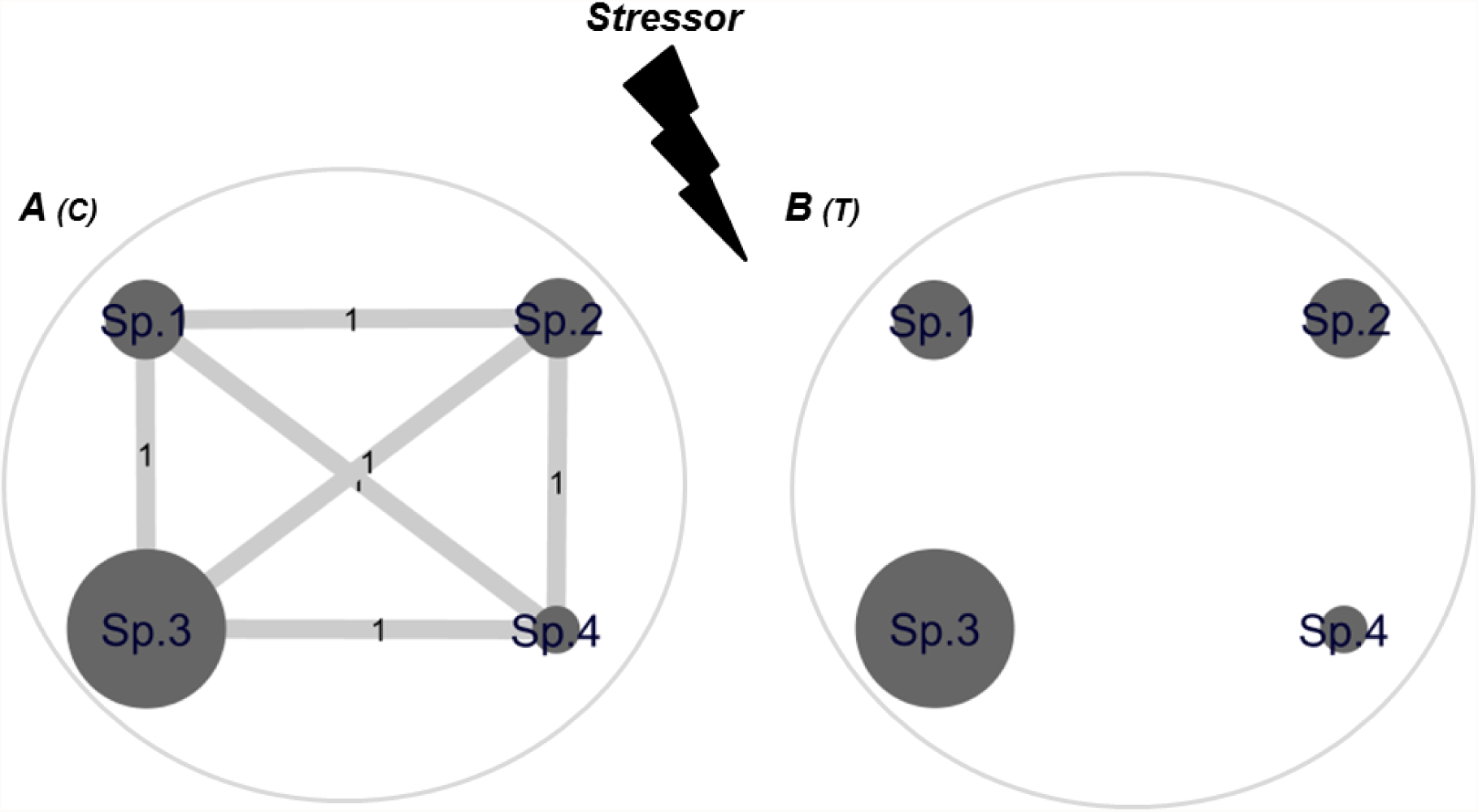
Hidden effects on the community caused by a stressor are exposed by the correlation network analysis. Significant Spearman correlation values are shown as edges (p□0.001), balls are related to relative specific abundances.

Changes in the relationship between specific fractions of a system after the presence of a stressor could affect the response of the community to a new stressor (Ashauer et al. 2017) by the modification of its resistance. In this sense, if the Sp.4 of our example were bloom-forming cyanobacteria, it could be possible that its disconnection from the community (Fig. III B) would enhance the possibility of bloom after a new stressor or opportunity (Fig. III C).

**Fig. III:**
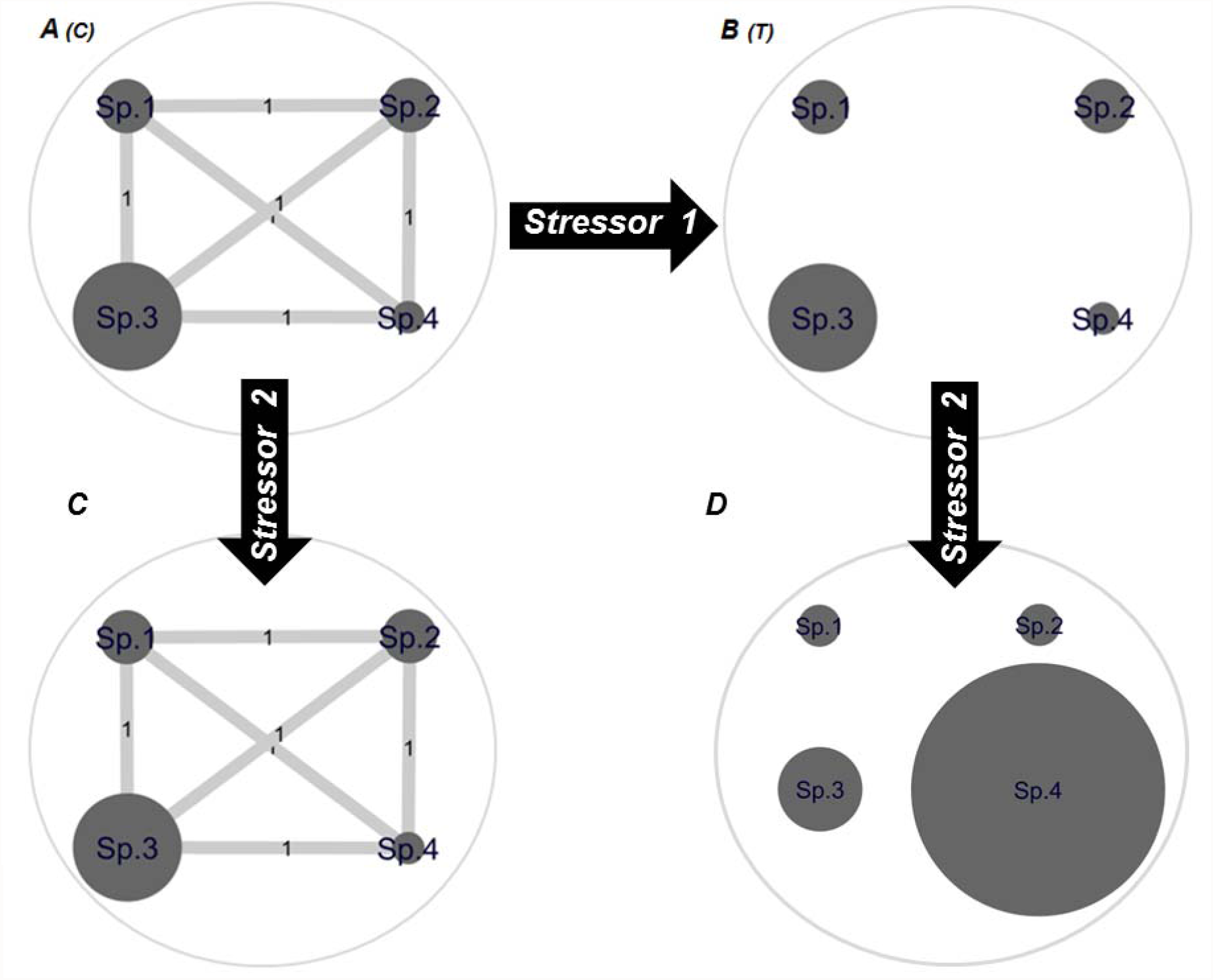
Possible consequences of changes in network behaviour facing sequential stressors.

We intend to motivate the incorporation of correlation networks analysis in the ecotoxicological assays that stand at the community level or higher. It is important to highlight that conclusions should be made paying attention to the inherent number of correlation possibilities, which is determined by the number of replicates and the number of components, as well as following a biological hypothesis. Correlation values must be calculated after normality verification. If this were the case, Pearson coefficients should be used, in other cases Spearman rank approach is recommended. Moreover, multi-trophic studies analysis could also be improved with this kind of approach (Fig. IV).

**Fig. IV:**
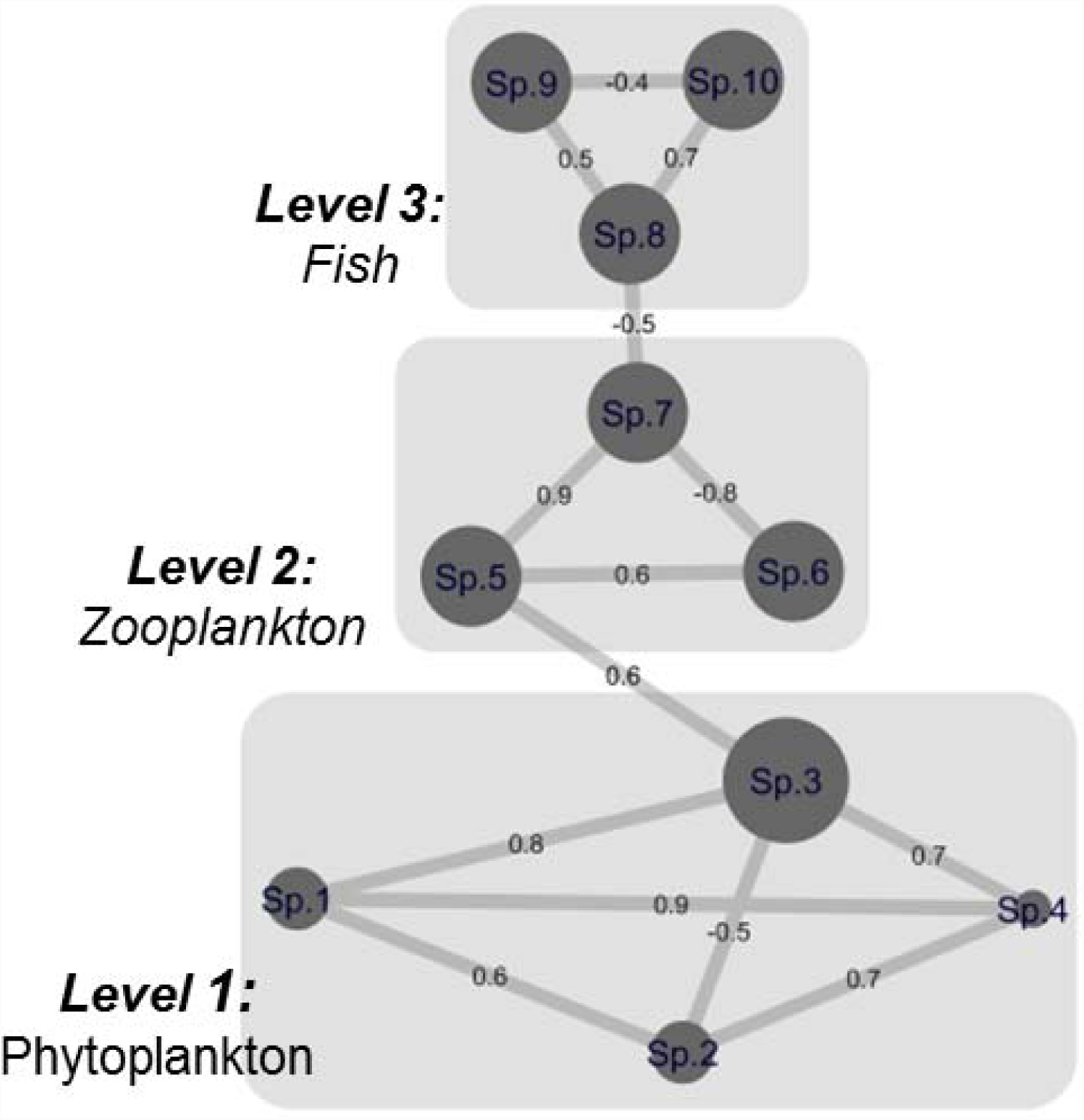
Multi-trophic correlation network.

## Acknowledgments

I thank my director Dr. Haydée N. Pizarro for stimulating my own ideas. This work was supported by PICT 2014.1586, UBACyT 20020130100248BA and PIP 11220130100399. The author declares that she has no conflict of interest. This article does not contain any studies with human participants or animals performed by any of the authors.

